# *rec-1* loss of function is insufficient to homogenize crossover distribution in *Caenorhabditis elegans*

**DOI:** 10.1101/2023.07.18.549456

**Authors:** Tom Parée, Luke Noble, João Ferreira Gonçalves, Henrique Teotónio

## Abstract

Meiotic control of crossover (CO) position is critical for proper homologous chromosome segregation and organismal fertility, recombination of parental genotypes, and the generation of novel genetic combinations. We here characterize the recombination rate landscape of a loss of function genetic modifier of CO position in *Caernorhabditis elegans*. By averaging CO position across hermaphrodite and male meioses and by genotyping 203 single-nucleotide variants covering about 95% of the genome, we find that the characteristic chromosomal arm-center recombination rate domain structure is lost in a loss of function *rec-1* mutant. The *rec-1* loss of function mutant smooths the recombination rate landscape but is insufficient to eliminate the non-uniform position of CO. We further find that the *rec-1* mutant is of little consequence for organismal fertility and embryo hatchability and thus for rates of autosomal non-disjunction. However, it specifically increases X chromosome non-disjunction rates and males’ appearance. Our findings question the maintenance of genetic diversity among *C. elegans* natural populations, and they further suggest that manipulating genetic modifiers of CO position will help map quantitative trait loci in low-recombining genomic regions.

## Introduction

Processes governing the number and position of crossovers (CO) during meiosis are tightly controlled, ensuring the segregation of homologous chromosomes into gametes and recombination of parental genotypes (Gray and Cohen 2016). Important variation of CO distribution patterns have been identified among and within species and can result from genetic modifiers (Chelysheva *et al*. 2012; Stapley *et al*. 2017; Brand *et al*. 2018; Brazier and Glémin 2022; Otto and Payseur 2019). Often, modifiers tend to affect the frequency of non-disjunction of homologous chromosomes, and thus gamete viability and organismal fertility, in addition to the frequency of new genetic combinations (Hodgkin *et al*. 1979; Koehler *et al*. 1996; Hillers *et al*. 2018; Holsclaw and Sekelsky 2021). For example, the 8-fold increase in genome-wide recombination rates seen in the mustard *Arabidopsis thaliana* double loss of function mutant *figl1/recq4* in an hybrid genetic background is accompanied by a ∼ 20% decrease in pollen viability, and no change in female fertility (Fernandes *et al*. 2018). In contrast, loss of function *xnd-1* alleles in the nematode *Caernorhabditis elegans* increase recombination rates only in gene-rich regions of autosomal chromosomes, at a 35% cost in embryonic survival (Wagner *et al*. 2010). Characterizing genetic modifiers of CO number and position is thus important to explain the heritability of recombination rate variation and the maintenance of genetic diversity in natural populations (Chelysheva *et al*. 2012; Stapley *et al*. 2017; Ritz *et al*. 2017; Dapper and Payseur 2017; Brand *et al*. 2018; Samuk *et al*. 2020; Lee *et al*. 2021). Further, their manipulation could be useful for breeding programs and for finding quantitative trait loci (QTL) located in low-recombining genomic regions normally refractory to discovery (Mieulet *et al*. 2018; Fayos *et al*. 2022).

The androdioecious nematode *C. elegans* has 6 holocentric chromosomes, one of which is the sex-determining chromosome – hermaphrodites are *XX* and males *X*∅. Most hermaphrodite and male meioses exhibit one CO per pair of homologous chromosomes, i.e., there is near complete CO interference, with a lower CO probability in central chromosomal domains than in the flanking chromosomal arms (Barnes *et al*. 1995; Hillers *et al*. 2018). This non-uniform CO pattern results in heterogeneous recombination rate “landscapes” across centers and arms, particularly in autosomes (Rockman and Kruglyak 2009; Bernstein and Rockman 2016), in common with other nematode species (Ross *et al*. 2011; Noble *et al*. 2021b; Teterina *et al*. 2022; Rillo-Bohn *et al*. 2021; Thomas *et al*. 2015).

Little is still known about the molecular causes of these heterogeneous recombination rate landscapes (Saito *et al*. 2013; Mets and Meyer 2009; Gerstein *et al*. 2010; Hillers and Villeneuve 2003; Henzel *et al*. 2011; Kaur and Rockman 2014; Wagner *et al*. 2010; Meneely *et al*. 2012; Hillers *et al*. 2018; Lascarez-Lagunas *et al*. 2023).

The central chromosomal domains encompass megabases of DNA sequence. They are characterized by higher gene density, open chromatin and highly expressed genes, and a greater proportion of highly conserved genes with essential functions on average (Barnes *et al*. 1995; Kamath *et al*. 2003; Parkinson *et al*. 2004; Gerstein *et al*. 2010; Liu *et al*. 2011). Central domains also show lower genetic diversity in wild populations than arms, due to the increased efficacy of linked selection (Cutter *et al*. 2009; Andersen *et al*. 2012; Rockman *et al*. 2010; Lee *et al*. 2021). Not unexpectedly, most QTL for morphological, physiological, and life-history traits have been found in the chromosomal arm domains; and when found in the centers QTL intervals can span hundreds to thousands of genes, e.g., (Andersen and Rockman 2022; Noble *et al*. 2017; Zhang *et al*. 2021, 2022; Vigne *et al*. 2021).

Several genes are known to play a role in the placement and number of CO in *C. elegans* (Zetka and Rose 1995; Meneely *et al*. 2012; Wagner *et al*. 2010; Youds *et al*. 2010; Saito *et al*. 2012; Hillers *et al*. 2018). Most of them, such as *xnd-1, him-5*, and *rtel-1* are associated with chromosome non-disjunction and increased embryonic lethality (Meneely *et al*. 2012; Wagner *et al*. 2010; Barber *et al*. 2008). The severity of non-disjunction differs across chromosomes, with *him-5* and *xnd-1* primarily affecting the X chromosome, and thus the appearance of males (Meneely *et al*. 2012; Wagner *et al*. 2010). One of the first recombination rate modifiers described for any organism was *rec-1* (Rose and Baillie 1979) identified by the appearance in a lab strain of a mutation that increased recombination in the center domains of chromosomes I, IV, and V (Rose and Baillie 1979; Rattray and Rose 1988). Later work showed that *rec-1* changes CO position, not CO number, and is associated with only a weak X chromosome non-disjunction phenotype and no apparent embryonic lethality (Rattray and Rose 1988; Zetka and Rose 1995; Chung *et al*. 2015). *rec-1* is located on the left arm of chromosome I and is known to promote double-strand break formation and resolution at prophase I of meiosis, sharing a partially redundant function in CO positioning with its paralog *him-5* (Chung *et al*. 2015).

The absence of a strong non-disjunction phenotype makes the *rec-1* interesting for studying CO patterning and the potential effects of *rec-1* on genetic diversity found in natural populations. One particular problem to understand is whether loss of function in *rec-1* is sufficient to eliminate the non-uniform placement of COs, and hence the heterogeneity of recombination landscapes along chromosomes. Current genetic linkage maps for loss of function *rec-1* mutants are based on a handful of markers across the *C. elegans* genome (Zetka and Rose 1995; Chung *et al*. 2015). Here, we report the construction of recombinant inbred advanced intercross lines (RIAIL) for *rec-1* wild-type and loss of function mutant alleles in different genetic backgrounds. By averaging CO position across hermaphrodite and male meioses, and by genotyping 203 SNV (single-nucleotide variants) covering 94.7% of the physical length of chromosomes, we find that the distinct arm-center recombination rate landscape is lost in the *rec-1* mutant. We demonstrate, however, that *rec-1* loss of function is insufficient to homogenize CO placement along chromosomes. We further confirm, across two genetic backgrounds, that the *rec-1* mutant is of little consequence for organismal fertility and thus for autosomal non-disjunction. As expected, rates of X chromosome non-disjunction are higher in *rec-1* mutants.

## Materials and methods

### *C. elegans* culturing

Strains obtained from the Caenorhabditis Genetics Center (University of Minnesota), or created by us, were stored at -80°C until usage (Stiernagle *et al*. 1999). Once thawed, samples were kept at 20°C and 80% RH in incubators and cultured in 6 cm or 9 cm Petri plates filled with NGM-lite agar (US Biological) and, respectively, a 10 uL patch or 100 uL lawn of *Escherichia coli* (HT115) cultured overnight. Synchronized cultures of starved L1 larvae were obtained by “bleaching” adults with a 20 mM KOH : 0.6% NaClO solution, with embryos kept in M9 solution at 20°C with aeration for 24h (Stiernagle *et al*. 1999). Hermaphrodites were visually distinguished from males and hand-picked or counted at the larval L4 or young adults stages under Nikon SM1500 dissecting microscopes.

### Parental RIAIL strains

For Recombinant Inbred Advanced Intercross Line (RIAIL) construction, we crossed six strains with three different genetic backgrounds (Table S1). The three *rec-1* wild-type allele strains used were EEV1401, EEV1402, and N2. The three *rec-1* loss of function mutant allele strains used were EEV1403, EEV1404, and BC313. EEV1401 was derived from 13 generations of self-fertilization of hermaphrodites from a polymorphic domesticated population, as described in Chelo *et al*. (2013b). Likewise, EEV1402 was derived by self-fertilization of hermaphrodites from a polymorphic lab domesticated population with an introgressed *ccIs4251* (*myo3p::GFP(NLS)::LacZ transgene*) construct, located in the center of chromosome I (Fire *et al*. 1998; Chelo *et al*. 2013b). This strain was used to tag the *rec-1* mutant with a morphologically visible marker for downstream analyses. N2 is the reference strain of *C. elegans* (WBStrain00000001). Mutant strain BC313 harbors the *rec-1(s180)* allele in an N2 background, as described in Rose and Baillie (1979). EEV1403 [*rec-1(eeg1403)*] and EEV1404 [*rec-1(eeg1404)*], two new *rec-1* mutant strains, were generated by directed mutagenesis in EEV1401 and EEV1402, respectively (see next).

### CRISPR-Cas9

CRISPR-Cas9-directed mutagenesis was used to generate an out-of-frame deletion in exon 2 of the *rec-1* gene and create a phenocopy of the *rec-1(s180)* allele (Chung *et al*. 2015). The target sequence (5’-GAACTGGATAACTGGCCGGC-3’; ordered from IDT) of the single guide RNA (sgRNA) was retrieved from Chung *et al*. (2015), and previously described CRISPR-Cas9 protocols were followed (Paix *et al*. 2017; Dokshin *et al*. 2018). Briefly, the sgRNA was generated by incubating 115 µM of trans-activating RNA (tracrRNA; IDT) and 85 µM of target RNA for 3 minutes at 95°C, followed by a decrease in temperature of 5°C every minute until 25°C. The final microinjection mixture was composed of recombinant *Streptococcus pyogenes* Cas9 nuclease V3 (IDT), 0.3 µg/µL, 1:3 of *rec-1* gRNA mix, 1:10 of *dpy-10* sgRNA mix, and 110 ng/µL of *dpy-10* repair template (Eurofins). This mix was incubated for 30 minutes at 37°C to form ribonucleoprotein complexes and injected into hermaphrodite adult gonads using a Transjector 5246 (Eppendorf). *F*_1_ progeny were singled out from plates displaying a high proportion of the dumpy (*dpy-10*) phenotype. After two days, the *F*_1_ were screened by PCR for the *rec-1* deletion at the expected site using an *HpaII* restriction enzyme (Thermofisher). PCR products harboring a deletion were Sanger sequenced at Eurofins, and an 8-bp deletion was selected (see Figure S1). *F*_2_ hermaphrodites of this genotype were backcrossed to males of their respective parental strains (EEV1401, EEV1402) for three generations.

### RIAIL construction

We used the Random Mating with Equal Contributions - Recombinant Inbred Advanced Intercross Line (RPMEC-RIAIL) design of Rockman and Kruglyak (2008) which allows for random mating with no variance in offspring number during the intercross stage. This design maximizes genetic linkage map precision while minimizing drift and selection. In detail (Figure S2), after mating between hermaphrodites of a parental strain and males of a different strain (P0), the *F*_1_*s* were sib-mated, and resulting *F*_2_*s* placed into 55 different 6 cm Petri dishes, each containing two males and one hermaphrodite. After that, and until *F*_5_, one hermaphrodite from each plate was randomly mated with two males from another plate. This is the intercross stage of RIAIL construction, though we use the term intercross for the whole protocol. Finally, and for the following 8 generations, we derived one inbred lineage from each plate by randomly picking a hermaphrodite and allowing it to self (Figure S2).

Wild-type *rec-1* RIAIL panels were generated by crossing EEV1401 with N2 or EEV1402, while mutant *rec-1* RIAIL panels were generated by crossing EEV1403 with BC313 or EEV1404 (Table S1). EEV1401 x N2 and EEV1403 x BC313 are referred to as type 1 intercrosses, while EEV1401 x EEV1402 and EEV1403 x EEV1404 are referred to as type 2 intercrosses (Table S2). Reciprocal crosses at *P*_0_ were done for each intercross. This resulted in eight *F*_3_ families for 420 individuals, from which we recovered 342 RIAILs, as some lines were lost during inbreeding (Table S2).

### DNA extraction

Starved nematodes from each line were harvested into 15 mL tubes with M9 isotonic solution, centrifuged at 8 rcf for one minute, with the pellet being re-suspended in M9 to remove remaining *E. coli*. Genomic DNA (gDNA) was extracted from each sample using the protocol described in Sunnucks and Hales (1996). Nematodes were lysed in TNES buffer, proteins precipitated at 15,000 rcf in 5M NaCl, and DNA precipitated at 15,000 rcf by adding pure ethanol. The DNA pellets were washed in ethanol 70% and air-dried before being resuspended in 50uL of TE buffer.

### RIAIL and parental strain genome sequencing

To genotype the RIAILs and parental strains, we used 2b-RAD sequencing (Wang *et al*. 2012). The 2b-Rad libraries were prepared using the same protocols, adapters, primers, and other reagents of Richaud *et al*. (2018). Briefly, we used the *Bcg1* restriction enzyme to cut 36 bp DNA fragments around its recognition site, which were then ligated with eight barcoded adaptors using T4 DNA ligase (NEB). The constructs were then amplified by PCR using sets of primers with an additional 24 barcodes, for 192 unique barcode combinations. The pooled samples were purified on a 2% agarose gel using a Nucleospin Macherey Nagel kit.

DNA samples of 331 RIAILs and 5 DNA samples from each parental strain were prepared in two libraries of 192 and 191 samples each (22 RIAIL were present in both libraries). The libraries were sequenced at the I2BC platform (Gif-sur-Yvette, France) with 51 bp single end reads on a *NextSeq 550* instrument (*NextSeq 500/550 High Output Kit v2, Illumina*). In total, 322 RIAIL were successfully sequenced.

We also whole genome sequences parental strains as some missing genotypes were present in the 2b-RAD sequences. Library preparation and whole-genome sequencing of EEV1401 and EEV1402 parental strains to an average 10X coverage was carried out by BGI (Hong-Kong).

### SNV calling and quality control

The 2b-Rad sequencing data were first demultiplexed using *process_radtag* in *STACKS* (Catchen *et al*. 2013), then filtered for the presence of the *Bcg1* restriction site and the primer W nucleotides, and trimmed to 36 bp using a custom *R* script (see trimming.R on GitHub repository).

All sequencing data were aligned to the WS245 *C. elegans* reference genome (downloaded from *Wormbase*, Harris *et al*. (2014)) using *BWA* (Li and Durbin 2009), allowing for two mismatches.

25,732 SNV were called with *freebayes* (Garrison and Marth 2012). The first filtering step involved checking for the presence of the reference and alternative alleles in at least ten RIAILs, or the presence of the SNV measured before in the lab domesticated population from which EEV1401 and EEV1402 were derived (Noble *et al*. 2017, 2021a). Congruence with the parental genotype was also confirmed. After this filtering step, the number of SNV that were kept dropped to 258. 209 markers segregate between the parental strains in at least one cross. At a subsequent filtering step, any SNV showing a higher pairwise LOD score with distant SNV, when compared with LOD scores with closer ones, was eliminated (Broman *et al*. 2003). 102 and 96 markers remained for type 1 and type 2 intercrosses, respectively, of which 56 were shared. On average, polymorphic SNV markers are found in 75% of the RIAILs, and, on average, for each RIAIL 94 markers were genotyped.

### MALDI-TOF genotyping

2b-RAD sequencing produced a highly uneven distribution of SNV. To increase SNV marker densities where needed, we genotyped 74 SNV from Noble *et al*. (2017) in two multiplexes using MALDITOF technology (Storm and Darnhofer-Patel 2003; Bradic *et al*. 2011) at the PGTB platform (Université de Bordeaux, Inrae). 61 SNV were successfully genotyped at 98% of the RIAILs. Their positions are indicated by orange triangles in Figure 1. All 61 SNV are polymorphic in type 1 intercrosses and only 51 in type 2 intercrosses. MALDI-TOF genotyping was done for all the 2b-Rad sequenced RIAIL and 14 additional RIAIL, thus totaling 336.

**Figure 1.**
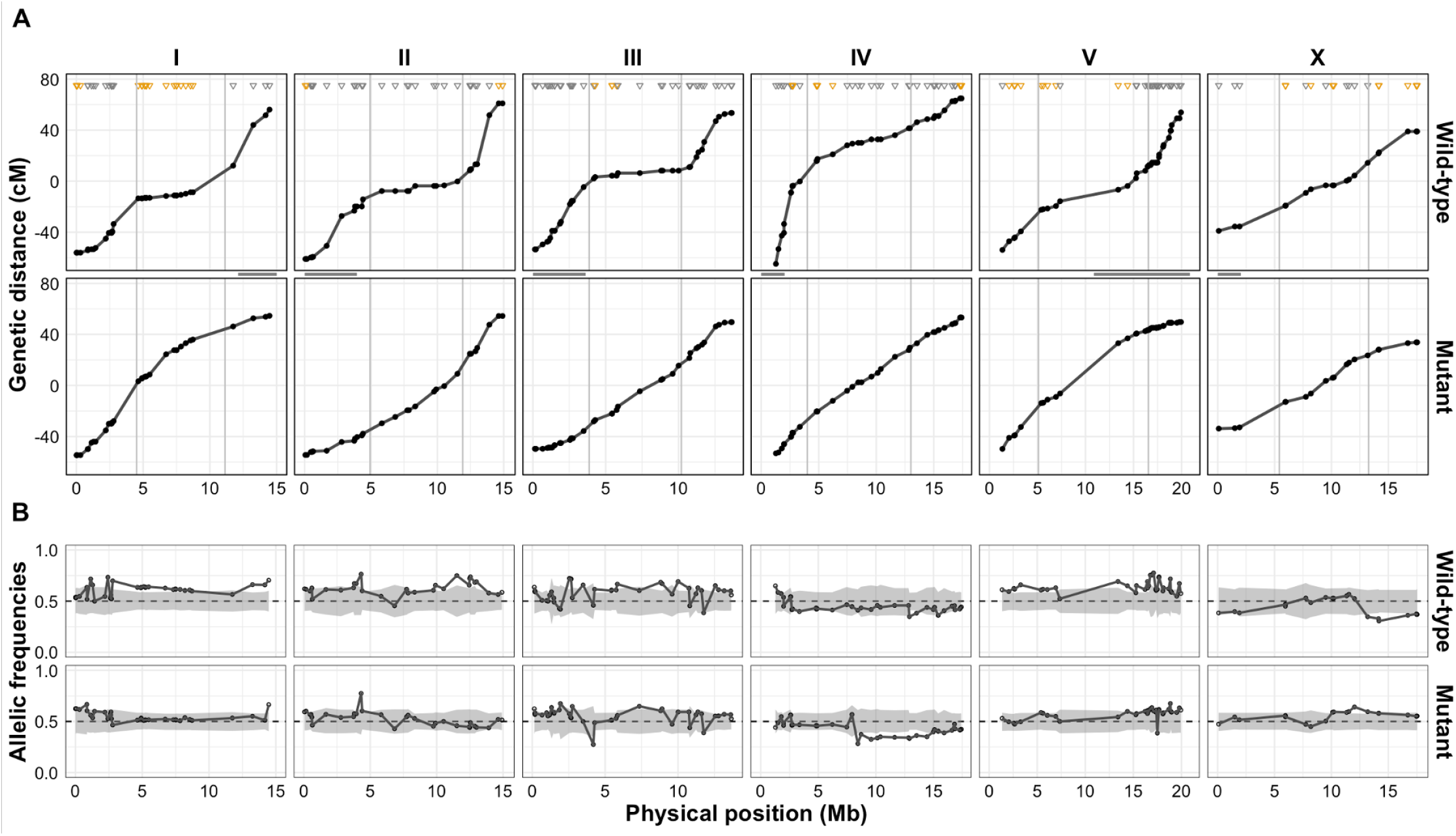
SNV genetic linkage among RIAILs. A. Consensus Marey plots showing the estimated genetic position (cM) of each SNV (single nucleotide variants) marker as a function of its physical position in the reference strain WBStrain00000001 (Mb). The top panels for each of the 6 chromosomes are from *rec-1* wild-type allele RIAILs, the bottom panels are from *rec-1* mutant allele RIAILs. The genetic position is centered in the middle. The vertical grey lines indicate the boundaries of the center domain as defined by Rockman and Kruglyak (2009). Grey and orange triangles indicate the markers obtained by 2b-RAD sequencing and MALDI-TOF genotyping. Horizontal grey lines between panels illustrate the location of the pairing centers (MacQueen *et al*. 2005). SNV encompass 94.7% of the genome (chr.I: 95.9%, chr.II: 97.2%, chr.III: 98.1%, chr.IV: 92.2%, chr.V: 88.8%, chr.X: 98.4%) B. Observed SNV allele frequencies polarized towards the EEV1401 parent allele among pooled *rec-1* wild-type (top) or mutant (bottom) RIALs. The shaded area is the 95% credible interval of expected allele frequencies obtained from 1,000 simulations of genetic drift and experimental sampling during RIAIL construction and genotyping

### Genetic linkage maps

The genetic linkage maps were constructed using the *est*.*map* function of the *R/qtl* package, using Haldane’s mapping function (Broman *et al*. 2003). RIAILs from reciprocal crosses were pooled, and the two intercross-specific maps were constructed for *rec-1* wild-type and mutant separately (Figure S4). Consensus maps were generated for both *rec-1* alleles, separately, without accounting for the type of intercross (Figure 1). Note that some genetic distances were estimated indirectly for consensus maps because SNV were not shared between the two types of intercrosses. We employ Marey plots for representation with the estimated SNV genetic position being plotted as a function of the reference genome physical position (WS245).

Eight RIAIL subpanels originated from 8 independent crosses varying for the intercross type, the *rec-1* allele, and the direction of the cross (see above). When pooled, the subpanel structure can bias the genetic linkage map estimation when strong allele frequency deviation from the expected 50% occurs differently in the different subpanels. An extreme case would be an allele nearly fixed in one panel and nearly lost in a second subpanel. Low heterozygosity in each panel also results in unobserved recombination events. Further, when pooling the two subpanels, imbalanced deviations can be masked and thus not accounted for in the LOD score calculation. To avoid these problems, we calculated the allele frequency deviation for each subpanel at each chromosome (Figure S3). The three subpanels with the higher average deviation were one wild-type subpanel for the V chromosome and two wild-type subpanels for the X chromosomes (one of each intercross type). Withdrawal of relevant data did not affect the chromosome V estimates but visibly changed X chromosome estimates (Figure S5). We thus discarded the data at the X chromosome from the two problematic subpanels for subsequent analysis.

### Simulations of drift and selection

We found the expected SNV allele frequency change from random sampling by running 1,000 individually-based simulations of the RIAIL construction with the realized individual numbers, family sizes, and missing genotypes (Figure 1). During the intercross stage of RIAIL construction, each hermaphrodite was randomly mated with one or two males from the same family. Offspring genotypes resulted from randomly picking one hermaphrodite gamete and one male gamete. During gametogenesis, one obligate CO per pair of the 6 homologous chromosomes was randomly placed along the chromosome with the placement probability depending on the wild-type or mutant *rec-1* consensus genetic distance between observed SNV. After fertilization, all offspring survive to adulthood except for random extinction mimiching the loss of lineage during the selfing phase. These simulations thus model genetic drift and experimental sampling during RIAIL construction and genotyping.

We further modeled viability selection during the RIAIL construction. For this, we assumed that offspring fitness is equal to 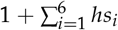, where *h* is a dominance coefficient (set to 0.5 in hermaphrodites and in male autosomes), and *s* a coefficient of the selected allele in chromosome *i*. Each simulation was set up with selected SNV alleles in only one of the parental strains. One random SNV at each chromosome was chosen with *s* = − 0.15. Thus, for example, when a given hermaphrodite was homozygous at six loci for the deleterious allele, it had a survival probability of only 0.1. Likewise, when a male was homozygous for the deleterious allele in the 5 autosomes and hemizygous for the deleterious allele in the X chromosome. The *R* script used for the simulations can be found in our GitHub repository.

### Genetic map differences between *rec-1* alleles

The Gini coefficient measures the inequity of the distribution of a variable and was used here to measure the heterogeneity of recombination rate landscapes. The Gini coefficient was calculated as in Kaur and Rockman (2014) by generating the Lorenz curves for each chromosome (Lorenz 1905) plotting the cumulative physical size of SNV intervals sorted by their recombination rates against their cumulative genetic size. The Gini coefficient is 2 times the area between the Lorenz curves and a uniform distribution (y=x), estimated using the trapezoidal approximation. Gini coefficients range from 0 (perfect equality of recombination rates along the chromosomes) to 1 (all recombination events occur at one locus). The *R* script used for estimation is in gini&AUC.R file in our GitHub repository.

To measure mean recombination rates per chromosomal domain, we used the domain boundaries of Rockman and Kruglyak (2009). For the left arm of the X chromosome, an adjacent SNV situated on the left of the center domain was used to better delimit the interval (see Table S4).

Null distributions were obtained by randomly permuting the *rec-1* wild-type and mutant RIAILs within intercross types. Consensus genetic linkage maps were constructed for 1,000 permuted datasets.

### Non-uniform crossover position

The curvature of the *rec-1* mutant consensus genetic linkage map was estimated by calculating the Area Under the Curve (AUC), as defined by the relative genetic position and the relative physical position. The AUC was estimated using the trapezoidal approximation. Deviations from 0.5 (*y* = *x*) indicate a non-uniform CO position. Null distributions were obtained as above for RIAIL construction under genetic drift and experimental sampling, assuming that the probabilities of CO positioning were a function of the physical distance between SNVs. The *R* script used for estimation can be found in the gini&AUC.R file on our GitHub repository.

### Chromosomal non-disjunction

While autosomal non-disjunction is lethal, X chromosome non-disjunction results in viable *XXX* hermaphrodites and *X*∅ males (Hodgkin *et al*. 1979). We measured hermaphrodite fertility and embryo hatching proportion as a surrogate for autosomal non-disjunction. For X chromosome non-disjunction rates we measured the number of males in the selfed-progeny of hermaphrodites.

EEV1401, EEV1402, EEV1403, and EEV1404 strains were thawed from stored samples and maintained in 9 cm Petri dishes for two generations under the “bleaching/hatch-off” protocol described above. In the third generation, starvation-synchronized *L*_1_ larvae were seeded at a density of 1,000 per 9 cm plate, and *L*_4_ staged immature hermaphrodites hand-picked 46h ±6h later to a 6 cm Petri dish with *E. coli*. These hermaphrodites were killed two days later, and fertility was calculated as the number of offspring reaching the adult stage three days later. Hatchability was calculated as the number of adult progeny relative to the number of embryos counted during the first day of embryo laying. The number of males was measured in the adult progeny of each hermaphrodite. The total progeny of 22,196 from 406 hermaphrodites was scored in 8 independent blocks, each corresponding to a different thaw and/or assay date. Hatchability was measured in a subset of 154 mothers in 6 blocks, for 8,480 embryos counted (Table S3).

Fertility, hatchability, and male numbers data was analysed with generalized linear models using the *glmer* function in *R*, accounting for the fixed effects of parental strain and the random block effects (*lme4* textitR package, Bates *et al*. 2015). Contrasts were done with the *emmeans* function of the *emmeans R* package (Lenth 2021). A Poisson distribution was assumed for fertility data, and a random effect for each observation was added to model overdispersion (Harrison 2014). Male numbers were modeled with a binomial distribution. For hatchability, we pooled counts for *rec-1* wild-type or *rec-1* mutant strains and performed a z-test. To compare the hatchability of the two different genetic backgrounds, we tested for differences in the slope of *adult*.*worms* ∼ 0 + *embryos*.

## Results

### Genetic maps are unaffected by genetic background

To study *rec-1* effects on CO positioning, we obtained recombinant inbred advanced intercross lines (RIAIL) from parental strains harboring either the *rec-1* wild type or a loss of function mutant alleles, in different genetic backgrounds (see Methods). Backgrounds-specific “type 1” intercrosses were generated between the N2 and EEV1401 parental strains for the wild-type RIAIL panel or between their corresponding background mutant strains BC313 and EEV1403 for the mutant panel (Table S2). Background-specific “type 2” intercrosses were generated between EEV1401 and EE1402, or EEV1403 and EEV1404, for the *rec-1* wild type and mutant panels, respectively. Each panel is further subdivided into two subpanels, only differing for the initial direction of the cross, thus totaling 8 subpanels (2 cross direction x 2 intercross types x 2 *rec-1* alleles). We estimated the number of CO events during RIAIL construction by genotyping single nucleotide variants (SNV) using a 2b-RAD sequencing approach combined with targeted MALDI-TOF genotyping to increase marker density and uniformity (see Methods). 149 *rec-1* wild-type and 187 *rec-1* mutant RIAILs were obtained (Table S2). After quality control, 163 SNV were retained in type 1 intercrosses and 147 SNV in type 2 intercrosses, with 107 markers shared between the two intercrosses, thus resulting in a total of 203 markers for analysis (see the GitHub repository).

Genetic linkage maps between SNV were estimated with the *R/qtl* package for the different intercross types and *rec-1* alleles, assuming a Haldane mapping function (see Methods). We compared the effect of the genetic background by using only shared SNV within intercross type and *rec-1* allele (4 different panels). Comparisons between intercrosses were then made at adjacent genetic intervals of at least 1 cM. We find that, among all intervals (45 wild-type and 52 mutant), only one in chromosome I differed between genetic backgrounds of the *rec-1* wild-type allele (Figure S4; P-Value=0.045, permutation-based with Bonferroni correction for multiple testing). This interval between I:7328409 and I:11744848 is shorter in type 2 intercrosses than in type 1 intercrosses. The GFP transgene segregating in the type 2 intercross panel at this location can explain these differences between genetic backgrounds. It was previously reported that the GFP transgene suppresses recombination (estimated genetic position: + 2.5 cM≈ 8.1 Mb) (Hsieh *et al*. 1999; Ceron *et al*. 2007). Other potential differences between genetic backgrounds, particularly in chromosome V (Figure S4), can be explained by overestimating genetic distances between physically distant SNV when using the Haldane mapping function, cf. (Broman *et al*. 2003). We can thus conclude that genetic linkage maps do not show differences between genetic backgrounds.

### *rec-1* consensus genetic linkage maps

By pooling RIAIL data across genetic backgrounds, we increased SNV numbers 1.3-fold and estimated consensus genetic linkage maps for the *rec-1* wild type and mutant alleles covering 94.7% of chromosomal physical lengths (Figure 1A).

At several locations across the genome, we find that observed SNV frequencies among RIAILs deviate from 50% (Figure 1B). We used simulations to determine whether these deviations were expected from experimental sampling and genetic drift during RIAIL construction. We simulated Mendelian SNV segregation, modeling CO position with the *rec-1* wild-type or mutant allele consensus genetic linkage maps, and using the sampling design of realized RIAIL construction and genotyping (see Methods). SNV allele frequencies depart significantly from neutral expectation for 50.2% of SNV for the *rec-1* wild-type allele panels, 24.6% for the *rec-1* mutant allele panels, and 61.1% for all RIAIL together. The simulations further suggest that the average absolute deviation from 0.5 is higher than expected by genetic drift and experimental sampling (P-value < 0.001; observed value different from the null distribution of 1,000 simulations).

SNV allele frequency deviations could result from residual self-fertilization in the parental generation (*P*_0_, Figure S2). We find little evidence of this because the average EEV1401 SNV allele frequency is higher when EEV1401 is the *P*_0_ hermaphrodites but it is not significant (57.3% compared to 54.1%; quasibinomial GLM: P-value=0.54). SNV allele frequency deviations could also result from selection/non-Mendelian segregation during the RIAIL construction. We find some evidence for selection as the EEV1401 strain allele (non-N2 and non-EEV1402) is more frequent among RIAILs (Figure 1B, Figure S3). Note, however, that this varies between chromosomes and that the EEV1401 allele is less frequent on chromosome IV.

As SNV allele frequencies among RIAILs depart significantly from neutral expectations, they could cause artefactual differences between the *rec-1* wild type and mutant allele consensus genetic linkage maps (Figure 1A). To address this issue, we first compared the total genetic map lengths to the expected total map length, per chromosome, from the simulations of RIAIL construction. For the observed consensus maps, the length of autosomes is longer than expected: there is a 1.42-fold total map expansion for the observed *rec-1* wild-type allele and a 1.24-fold map expansion for the observed *rec-1* mutant allele (comparison with null distribution from the simulations: P-value for *rec-1* wild-type, chr.I: 0.017; chr.II:0.009; chr.III:0.004; chr.IV: <0.001; chr.V: <0.001; chr.X: 0.047; genome: <0.001; P-value for *rec-1* mutant, chr.I: 0.028; chr.II:<0.001; chr.III:0.004; chr.IV: <0.001; chr.V: 0.14; chr.X: 0.143; genome: <0.003).

Because selection could also change estimated map length, we simulated RIAIL construction as above, but with strong viability selection against one of the parental strains (10-fold fitness reduction; see Methods). These simulations, however, show that selection on average does not change the map length when compared to drift (for genome, comparison of the genetic length distributions of 1,000 drift and 1,000 selection simulations for genome total genetic size: t-test: P-value = 0.47).

We next compared the *rec-1* wild-type allele consensus genetic maps to the reference genetic linkage maps of Rockman and Kruglyak (2009), which were obtained from a cross between N2 and CB4856 strains at much higher SNV densities. Both our maps and those of Rockman and Kruglyak (2009) are congruent (Figure S6), with a Pearson correlation between SNV genetic positions for each chromosome among the two genetic maps − 0.98.

### The *rec-1* mutant smooths the recombination rate landscape

Visual inspection of the Marey plots presented in Figure 1A indicates that the *rec-1* mutant allele smooths the recombination rate landscapes of all autosomes, and less so for the X chromosome. To quantify this smoothing, we compared the heterogeneity of recombination rates for all chromosomes between the wild-type and mutant consensus maps using Gini coefficients (see Methods). The *rec-1* mutant allele is expected to reduce the recombination rate differences between recombination chromosomal domains and thus lower Gini coefficient values. As expected, the *rec-1* mutant allele maps have lower Gini coefficients for all autosomes, though less so for chromosome I (Figure 2, Table S5). We used permutation tests, changing *rec-1* allele identity (see Methods), to compare Gini coefficients between the *rec-1* wild-type and mutants (insets in Figure 2). We find that observed values are significantly different to null distributions for chromosomes II, III, IV, and V (P-values: chr.II: <0.001, chr.III: <0.001, chr.IV: <0.001, chr.V: 0.002). For chromosomes I and X, there are no differences in Gini coefficients of mutant and wild-type maps (permutation P-value: chr.I: 0.21, chr.X: 0.84).

**Figure 2.**
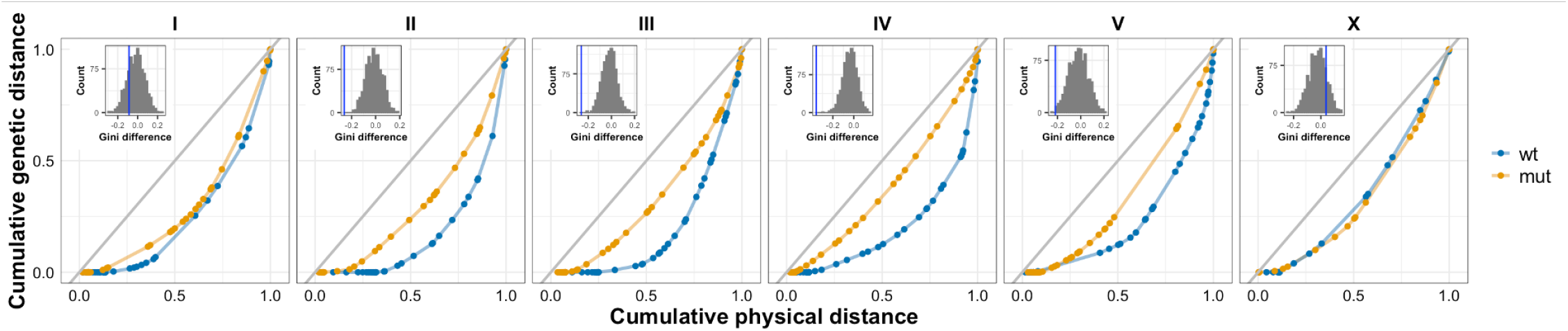
Heterogeneous recombination landscapes. Lorenz curves for each chromosome are shown for the *rec-1* wild-type (blue) and *rec-1* mutant (orange) alleles, generated by plotting the cumulative physical size of SNV intervals sorted by their recombination rates against their cumulative genetic size. The Gini coefficient is 2 times the area between the Lorenz curves and a uniform distribution of recombination rates (the grey line). Insets in each panel, for each chromosome, show the difference in Gini coefficient distribution of RIAIL permutations irrespective of *rec-1* allele identity; blue vertical line, the observed Gini coefficient difference between *rec-1* wild type and mutant alleles.

By eliminating the recombination rate chromosomal domain structure of the *rec-1* wild-type allele, the *rec-1* mutant allele, on average, decreases recombination in the arms and increases recombination in the center domains (Figure 3). The fold-change in average recombination rates per domain is significant for all intervals tested except in the left arm of I (IL), on the right arm of IV (IVR), on the left arm of V (LV), on the left arm of X, and in the center of chromosome X (Table S4). The asymmetry in the wild-type recombination landscape creates an inverse correlation of the mutant effect between arms (Pearson r = - 0.91; P-value = 0.01) and, in part, explains why the *rec-1* mutant does not affect IL, IVR, or LV; i.e., the *rec-1* mutant has a greater effect in the more recombinogenic arm than the less recombinogenic arm. Though the *rec-1* mutant has little effect on the total genetic length of the left arm of chromosome I, it affects the distribution of recombination by increasing it to the left of the subdistal regions, consistently with previous observations (Rattray and Rose 1988).

**Figure 3.**
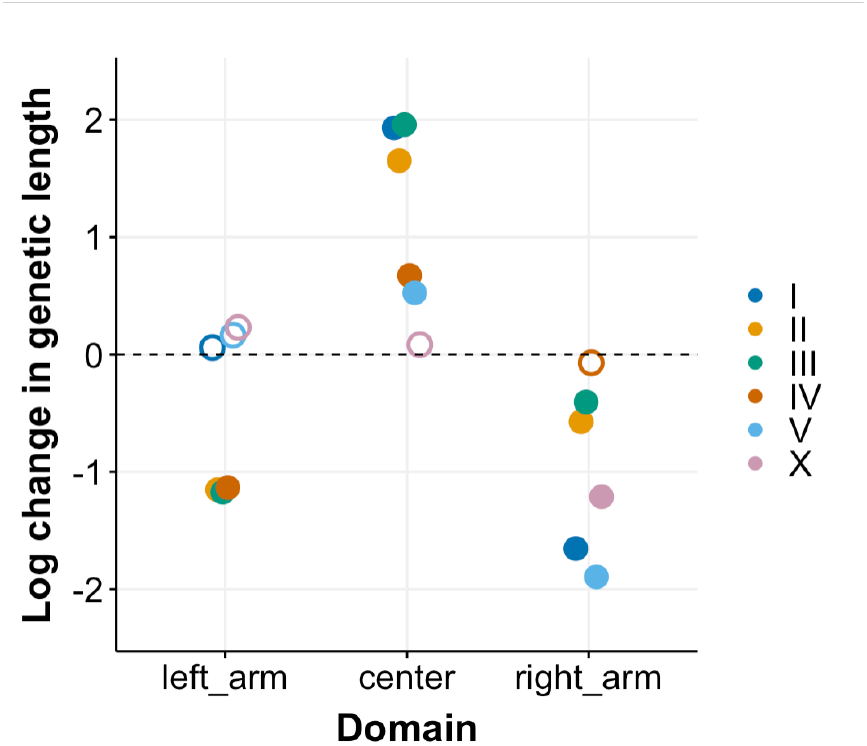
The *rec-1* mutant eliminates wild-type chromosomal recombination rate domains. The colored dots are the logarithm of the mutant and wild-type interval genetic length ratio for the several recombination chromosomal domains. The value is negative when the mutant decreases recombination rates in an interval and positive otherwise. Filled circles show significant differences from the null distribution, open circles non-significant, permutations of RIAIL *rec-1* allele identity. The chromosomal recombination rate domains were delimited as in Rockman and Kruglyak (2009). Interval values can be found in Table S4.

To test if the observed Gini coefficients for the *rec-1* mutant maps are consistent with a homogeneous recombination landscape, we used simulations as above but with CO position given only by the SNV physical position in the chromosome (see Methods). We find that the *rec-1* mutant has Gini coefficients consistent with a homogeneous recombination landscape only for chromosomes III and IV (Table S5).

Finally, we tested if *rec-1* impacts recombination at the chromosomal subtelomeric tip domains as defined by Rockman and Kruglyak (2009). For each mutant RIAIL, we counted the number of CO arising in each SNV interval spanning the tips that could be observed. We find that no CO was inferred for all examined intervals within the tips. Given the number of lines and the cumulative physical size of the examined intervals, if tips recombined at a genome-wide average rate, we would expect 15 CO among all RIAIL, which is much higher than the observed value (Chi-squared test: P-value = 9.7× 10^−5^). Hence, recombination is suppressed in the *rec-1* mutant at the chromosomal tips.

### The *rec-1* mutant is insufficient to eliminate non-uniform CO position

While the *rec-1* mutant smooths the *C. elegans* recombination landscape, significant heterogeneity in recombination rates remains, at least for chromosomes I, II, V, and X, and subtelomeric regions. To appreciate this heterogeneity we characterized in detail the curvature of the Marey plots shown in Figure 4. To estimate this curvature, we calculated the area under the relative genetic distance against the relative physical distance (see Methods). The curvature is different than expected for all chromosomes, except the X chromosome, with a homogenous recombination landscape (Figure 4). It should be noted that the relative area under the curves in the simulations with homogenous recombination landscapes is not centered on 0.5, particularly for chromosomes IV, and V, indicating that the genetic linkage map estimation algorithm induces a curvature despite the homogenous recombination rate landscape. This artifact is likely due to the uneven density of SNV together with problems in using the Haldane mapping function for physically distant SNV.

**Figure 4.**
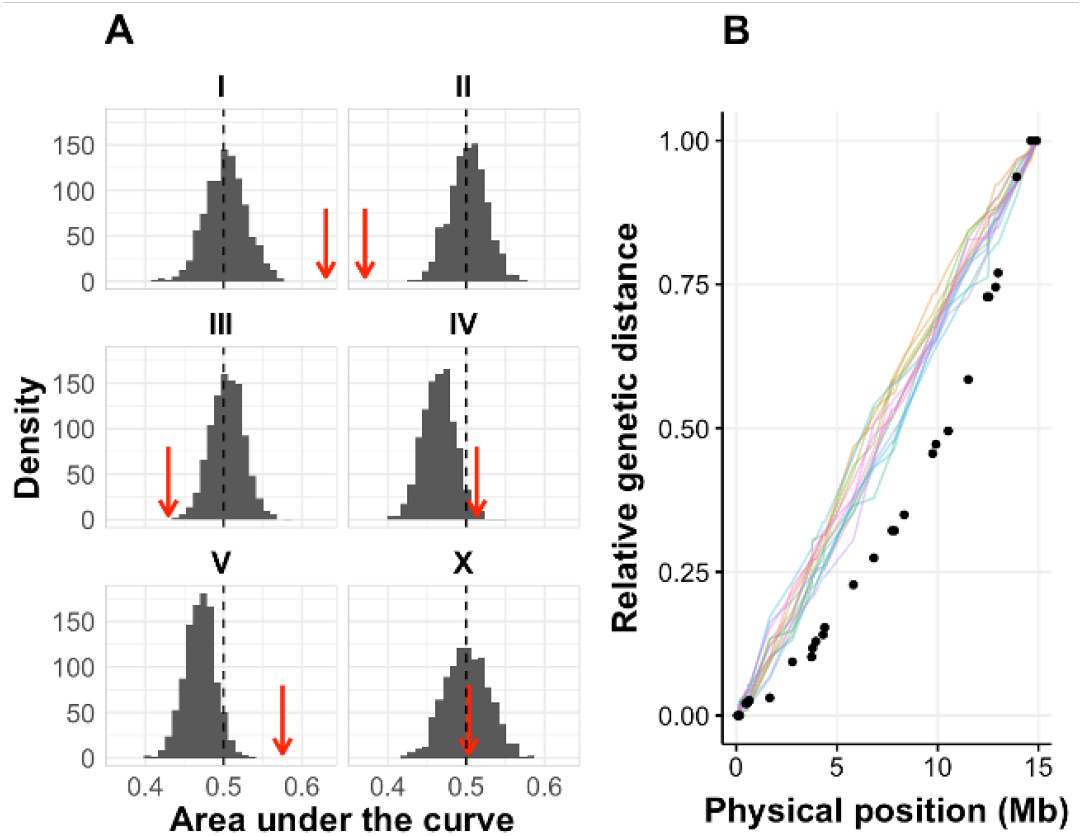
The *rec-1* mutant recombination landscape is not homogeneous. A. For each chromosome, the histogram (30 bins) of the relative area under the curve (AUC) of Marey plots was calculated in simulation with homogeneous recombination landscapes. The dotted black line indicates the relative AUC of 0.5 from a non-curved Marey plot. The red arrow indicates the observed relative AUC. B. The Marey plot for the observed genetic distances (black dots) in chromosome II is plotted as an example next to 15 randomly sampled simulated Marey plots with a random CO position and homogeneous recombination landscape (lines).

The potential artifact of the estimation algorithm is accounted for in simulations, but we further retrieved the curvature of genetic linkage maps for chromosome I and III from previously-published studies using a handful of markers in mutant *rec-1* (Zetka and Rose 1995; Chung *et al*. 2015). Genetic linkage maps were also retrieved for mutants *xnd-1* and *him-5* that are expected to functionally act upstream or redundantly with *rec-1* and to share similar pheno-types (Zetka and Rose 1995; Chung *et al*. 2015). We find that the curvatures of these previously-published genetic linkage maps are in the same direction as the *rec-1* linkage maps reported here (Figure S7). The results shown in Figure 4 should be, at least partially, revealing of heterogeneous *rec-1* mutant recombination landscapes.

### Chromosomal non-disjunction

Recombination modifiers are known to cause non-disjunction and embryonic lethality if on the autosomes, and increased male proportion in the progeny of selfed hermaphrodites if on the sex chromosome. We measured 96h post-L1 hermaphrodite fertility, hatchability, and male proportion in EEV1401 and EEV1402 genetic backgrounds with either of the *rec-1* alleles.

Fertility is not affected by the *rec-1* allele in either genetic background (Figure 5A). However, fertility is lower in the EEV1401 genetic background than in the EEV1402 background (GLM: P-value = 0.0017), consistently with (Chelo *et al*. 2013b).

**Figure 5.**
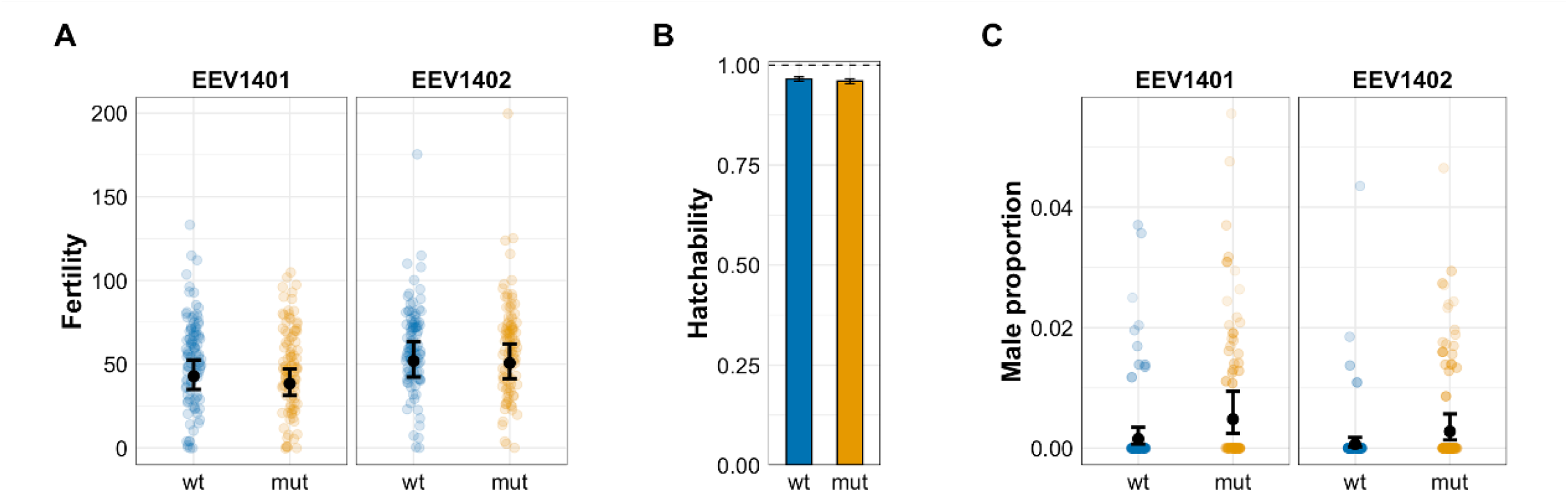
The *rec-1* does not impact homolog chromosome non-disjunction. Effects of wild-type (blue) and *rec-1* mutant (orange) alleles on fertility (A), hatchability (B, C), and male numbers (D) assayed in two different genetic backgrounds. The EEV1401 background corresponds to the EEV1401 (*rec-1* wild-type) and EEV1403 strains (*rec-1* mutant). The EEV1402 background corresponds to the EEV1402 (*rec-1* wild-type) and EEV1404 strains (*rec-1* mutant). A. Each colored dot is the 96h post-L1 fertility: the number of self-fertilized adult progeny of one hermaphrodite. The black dots and error bars are the mean and 95% CI estimated by a generalized linear model. B. Hatchability is the total number of adults divided by embryos laid. C. Colored dots are the number of males occurring among the progeny of selfed hermaphrodites. Opacity is determined by progeny number. Two dots corresponding to progeny under 30 are not shown. The black dots and error bars are the mean and 95% CI estimated by a generalized linear model.

Hatchability, measured as the number of viable adult progeny divided by the number of laid embryos, is not significantly different between the different *rec-1* alleles (z-test: p=0.20) (Figure 5B). Because some embryos were undetected, hatchability was in some cases above 100%. To compare the different strains’ hatchability, we tested for the difference between the slopes of hatched viable worms as a function of the initial number of embryos (see Methods). The EEV1401 genetic background has a lower hatchability than the EEV1402 genetic background (Figure S8).

Male numbers were 3.2 and 4.2-fold higher in the *rec-1* mutant compared to the *rec-1* wild-type for the EEV1401 and the EEV1402 genetic backgrounds (Figure 5C; GLM: P-value EEV1401: 0.0032; p-value EEV1402: 0.0079). The male proportion in the selfed progeny of hermaphrodites is about two times higher in EEV1401 than the EEV1402, regardless of *rec-1* alleles (GLM: genetic background effect, P-value = 0.034).

## Discussion

We have used the RPMEC-RIAIL design of Rockman and Kruglyak (2008) to generate high SNV (single-nucleotide variants) density consensus genetic linkage maps for *rec-1* wild type and loss of function mutant alleles (Figure 1). Our maps were constructed assuming complete crossover (CO) interference and were averaged over hermaphrodite and male meioses. The modeling of expected SNV (single-nucleotide variants) marker deviations from the random sampling during RIAIL construction and genotyping shows that *rec-1* wild type and loss of function mutant alleles do not bias genetic linkage map estimation. Our estimates show that, as expected, the wild-type recombination rate landscape is characterized by chromosomal central domains with less recombination than chromosomal arm domains (Barnes *et al*. 1995; Rockman and Kruglyak 2009). Besides the clear arm-center recombination rate domain structure at autosomes, known peculiarities are also observed with, for instance, a highly recombinogenic chromosome IV left arm. Recombination domain structure on the X chromosome was less obvious, though also, as expected, there were lower recombination rates in the middle of the chromosome (Rockman and Kruglyak 2009).

This wild-type recombination rate landscape is lost in the *rec-1* mutant (Figure 2,Figure 3) even if heterogeneity in crossover (CO) positioning is maintained (Figure 2, Figure 4). The *rec-1* mutant eliminates the arm-center domain structure (Zetka and Rose 1995; Chung *et al*. 2015), thus expanding genetic distances in the center domains and shrinking them in the arms domains. However, because neither the wild-type nor mutant recombination landscape is symmetric around the center domains, the change in recombination rate strongly varies depending on the region considered. Reduced recombination in *rec-1* mutant’ arm domains vary between only a 1.06-fold reduction for the left arm of chromosome I to a 6.6-fold reduction for the right arm of chromosome V (Figure 3). Interestingly, even though the average recombination rate in the left arm of chromosome I is unchanged, the *rec-1* mutant still increases recombination rates near the subtelomeric tip when compared to the *rec-1* wild-type allele (Zetka and Rose 1995; Rattray and Rose 1988). The *rec-1* mutant retains low recombination rates in the subtelomeric chromosomal tips, showing that different molecular pathways controlling CO position in these regions are maintained. It is known that the wild-type recombination landscape mirrors the distribution of double-strand breaks (DSBs) that initiate CO resolution during prophase I of meiosis. The exception is for the subtelomeric chromosomal tips, which have high DSB rates but show fewer COs, where *cis*-acting molecular factors may determine recombination suppression (Yu *et al*. 2016).

The *rec-1* mutant recombination rates gradually decrease or increase along the autosomes. This “curvature” of the genetic linkage map cannot be explained by genetic drift or experimental sampling during our RIAIL construction, and confirming our results, loss of function mutants in *him-5* and *xnd-1* also show a similar curvature on chromosomes I and III as the *rec-1* mutant studied here (Figure S7). The determinants of asymmetric recombination landscape in the rec-1 wild-type and mutant are unknown, though some patterns correlate with the pairing center position. For instance, the less-sharp boundary is on the side of the chromosome holding the pairing center for five chromosomes (Rockman and Kruglyak 2009). The arm of the chromosome holding the pairing center is associated with enriched in H3K9 methylation, a heterochromatin marker present in higher levels at the arms but depleted at the centers (Gerstein *et al*. 2010; Liu *et al*. 2011). The pairing center is always found on the left side of the chromosome except for chromosomes I and V, which are the only two autosomes with pronounced concave curvatures in the *rec-1* mutant Marey plot (see Figure 1; MacQueen *et al*. (2005); Phillips *et al*. (2009)). Hence, the less recombinogenic autosomal side and the position of the pairing centers generally correlate, the exception being chromosome IV.

The role of *rec-1* during meiosis remains poorly understood, though it shows striking similarities with the roles of *him-5* and *xnd-1*. Loss of function mutants of all three genes eliminate the recombination landscape and cause the non-disjunction of homologous chromosomes, particularly of the X chromosome. Our phenotypic assays for non-disjunction did not measure autosome non-disjunction and gamete viability as a mechanism to cope with aneuploid gametes (Vargas *et al*. 2017), and so it remains unclear whether *rec-1* mutant affects autosomal non-disjunction (Chung *et al*. 2015). The three genes, *him-5, xnd-1* and *rec-1* appear to be the central players of CO positioning by promoting double-strand breaks (DSB) formation because they all limit RAD51 deposition (a marker of DSB) and can be rescued by ionizing radiation causing DSB (Wagner *et al*. 2010; Meneely *et al*. 2012; Chung *et al*. 2015). The paralogous pair of *rec-1* and *him-5* have redundant functions as their double mutant leads to severe DSB deficiency (Chung *et al*. 2015). However, the *rec-1* mutant non-disjunction phenotype is much weaker than that of the *him-5* mutant. The *xnd-1* gene acts as a downstream factor of *him-5* and is thought to be a transcription factor (Yu *et al*. 2016). While *HIM5* and *XND1* proteins are located on the autosomes, they are mainly involved in DSB formation on the X chromosome, which may explain that loss of function mutants in these genes causes strong disjunction in the X chromosome (Wagner *et al*. 2010; Meneely *et al*. 2012). It is unclear, however, how DSB control on the X chromosome changes CO placement on the autosomes. One possibility is that an unknown meiotic checkpoint is activated, forcing more permissive CO placement. In line with this explanation, *xnd-1* accelerates RAD51 deposition after DSB formation, while *him-5* and *rec-1* mutants delay RAD51 deposition (Wagner *et al*. 2010; Meneely *et al*. 2012; Chung *et al*. 2015). Asynapsis of the X chromosome is sufficient to induce the delay in RAD51 deposition and impaired CO control on autosomes (Carlton *et al*. 2006). And while *him-5, xnd-1*, and *rec-1* might not be required for initial synapsis, loss of function mutants in these genes can cause desynapsis (Wagner *et al*. 2010; Meneely *et al*. 2012). Overall, *rec-1, him-5*, and *xnd-1* are required for DSB formation and possibly CO resolution, but they affect the segregation of homologous chromosomes differently.

Reduced recombination rate in the middle of the chromosomes is typical of many eukaryotes (Haenel *et al*. 2018; Brazier and Glémin 2022; Otto and Payseur 2019; Lenormand *et al*. 2016). Furthermore, the size of these low recombining regions positively correlates with overall chromosome size. This pattern could be due to COs occurring primarily at a fixed distance from the chromosome tips. For monocentric species, chromosome undergo a characteristic “bouquet” formation with telomere anchored to the nuclear periphery where CO-promoting factors may locate (Haenel *et al*. 2018; Brazier and Glémin 2022). At the end of prophase I of meiosis, the sister chromatids’ kinetochores face the same direction, while the homologs’ kinetochores face opposite directions resulting in the. Sister chromatids are maintained together by the centromere and when their cohesion is lost homologs can be properly segregated (Rubin *et al*. 2020). However, higher recombination rates in *C. elegans* chromosomal arms cannot be due to CO-promoting factors at the nuclear periphery as *C. elegans* holocentric chromosomes do not form bouquets, and only one chromosomal tip is anchored to the nuclear periphery by the pairing centers (Hillers *et al*. 2018; Melters *et al*. 2012; Mandrioli and Carlo Manicardi 2012). In the absence of localized centromeres maintaining sister chromatids, cohesion need to be partially kept to only separate homolog chromosomes (Melters *et al*. 2012; Mandrioli and Carlo Manicardi 2012). After the loss of sister chromatid cohesion, a single off-centered chromosome will form the cruciform-shaped bivalent with short and long arms resulting from asymmetric chiasma placement (Nabeshima *et al*. 2005; Martinez-Perez *et al*. 2008; Hillers *et al*. 2018). This asymmetry is essential for accurate reductional segregation because different factors are located in the short and long arms, factors that will inhibit the premature loss of cohesion between sister chromatids (De Carvalho *et al*. 2008; Tzur *et al*. 2012; Rogers *et al*. 2002). When a CO occurs in the middle of the chromosome, it will lead to symmetrical bivalents and premature sister separation (Altendorfer *et al*. 2020). It is thus puzzling that the *rec-1* mutant, increasing CO number in autosomal centers, does not cause substantial embryonic lethality or decreased organismal fertility (this study; Rose and Baillie 1979; Rattray and Rose 1988; Chung *et al*. 2015).

The fact that the *rec-1* mutant does not affect embryonic lethality or organismal fertility raises the question of whether *rec-1* polymorphism is relevant to genetic diversity among natural populations. Diversity in the coding regions of *rec-1* is present among natural populations (observed in CeNDR collection of wild isolate Cook *et al*. (2017)), though functional effects on recombination rate landscapes are unknown. This diversity could potentially be maintained via effects on X chromosome non-disjunction, independently of recombination, because males can be favored by natural selection in certain environmental conditions, e.g., (Teotónio *et al*. 2006; Teotónio *et al*. 2012; Morran *et al*. 2009a,b; Wegewitz *et al*. 2008). In this context, it is interesting to note that the *him-5* null mutant, which causes substantial X chromosome non-disjunction (though together with autosome non-disjunction), is a paralog of *rec-1* that presumably duplicated from an ancestral *rec-1*/*him-5* gene after the appearance of hermaphroditism from an ancestral male-female state (Chung *et al*. 2015). However, the arm-center chromosomal recombination rate domains structure is observed in several nematode species (Rockman and Kruglyak 2009; Ross *et al*. 2011; Noble *et al*. 2021b; Teterina *et al*. 2022; Rillo-Bohn *et al*. 2021; Thomas *et al*. 2015; Lee *et al*. 2021), and diversity at the *rec-1* locus may instead be maintained at mutation-drift equilibrium. In this situation, the evolution of genome composition could explain heterogeneous recombination rate landscapes. Different *Caenorhabditis spp*. show chromosomal recombination domains that correlate with gene density, the presence of operons, transposons, and chromatin features (Barnes *et al*. 1995; elegans Sequencing Consortium 1998; Blumenthal *et al*. 2002; Kamath *et al*. 2003; Gerstein *et al*. 2010; Liu *et al*. 2011). Even if *rec-1* increases recombination rates in chromosomal center domains, it may be of little consequence because natural populations are depauperate of genetic diversity in these regions (Cook *et al*. 2017; Crombie *et al*. 2019). When new allelic combinations do appear, they are likely selected against due to the presence of genotype incompatibilities (Dolgin *et al*. 2007; Chelo *et al*. 2013a; Seidel *et al*. 2008; Chelo and Teotónio 2013; Noble *et al*. 2021b). Characterizing the recombination rate landscapes and chromosomal non-disjunction patterns between parental *C. elegans* strains with higher genetic divergence in the central chromosomal domains than the ones used here, as it is now possible using highly divergent natural populations (Cook *et al*. 2017; Crombie *et al*. 2019; Lee *et al*. 2021), will help resolve these questions.

In conclusion, *rec-1* loss of function eliminates the characteristic landscapes of defined chromosomal recombination rate domains. However, the crossover position in loss of function *rec-1* continues to be non-uniform along the chromosomes. Future studies should determine the role of natural *rec-1* variation in CO positioning and chromosome non-disjunction to explain the genetic diversity found among natural populations of *C. elegans*.

## Data availability

Strains are available upon request. Data, R scripts, and modeling results can be found at our GitHub repository. Supplementary materials include six tables and eight figures.

## Acknowledgments

We thank V. Fernandes, H. Gendrot, A. Richaud, and R. Weissman for help with worm handling and molecular biology, and C. Dillmann, M.-A. Félix, T. Lenormand, F. Mallard, M. Rockman, D. Roze, A. Le Rouzic, and J. Yanowitz for discussion.

## Supplementary Tables

**Table S1.** Designation and genotypes of parental strains used for RIAIL construction.

| Strain | Background | rec-1 | GFP | reference |
| --- | --- | --- | --- | --- |
| N2 | N2 | wt | wt | Brenner (1974) |
| BC313 | N2 | *s180* | wt | Rose and Baillie (1979) |
| EEV1401 | A6140 | wt | wt | Chelo *et al.* (2013b) |
| EEV1402 | A6140GFP | wt | *ccls4251* | Chelo *et al.* (2013b), Fire *et al.* (1998) |
| EEV1403 | EEV1401 | *eeg1403* | wt | here |
| EEV1404 | EEV1402 | *eeg1404* | *ccls4251* | here |

**Table S2.** RPMEC-RIAIL subpanels’ designations and sample sizes during construction and genotyping.

| Cross | Hermaphrodite P0 | Male P0 | rec-1 | Intercross type | n in F2 | n in F13 | 2b-Rad | MALDI-TOF |
| --- | --- | --- | --- | --- | --- | --- | --- | --- |
| A | EEV1401 | N2 | Wild type | 1 | 55 | 33 | 29 | 33 |
| B | N2 | EEV1401 | Wild type | 1 | 55 | 37 | 35 | 37 |
| C | EEV1401 | EEV1402 | Wild type | 2 | 55 | 39 | 38 | 38 |
| D | EEV1402 | EEV1401 | Wild type | 2 | 55 | 41 | 40 | 41 |
| E | EEV1403 | BC313 | Mutant | 1 | 55 | 49 | 47 | 49 |
| F | BC313 | EEV1403 | Mutant | 1 | 55 | 44 | 41 | 43 |
| G | EEV1403 | EEV1404 | Mutant | 2 | 55 | 51 | 45 | 47 |
| H | EEV1404 | EEV1403 | Mutant | 2 | 55 | 48 | 47 | 48 |
| **Total:** |  |  |  |  | 440 | 342 | 322 | 336 |

**Table S3.** Number of hermaphrodites assayed for chromosomal non-disjunction and total brood size. Block refers to different parental strain thaw or assay dates. Indicator variables for which blocks were measured for fertility, hatchability and male numbers.

| Block | Mothers | 96h-progenies | Fertility | Spontaneous males | Hatching |
| --- | --- | --- | --- | --- | --- |
| 1 | 20 | 851 | yes | yes | yes |
| 2 | 20 | 830 | yes | yes | yes |
| 3 | 20 | 993 | yes | yes | yes |
| 4 | 31 | 2100 | yes | yes | yes |
| 5 | 32 | 2185 | yes | yes | yes |
| 6 | 31 | 1205 | yes | yes | yes |
| 7 | 108 | 4825 | yes | yes | no |
| 8 | 144 | 9207 | yes | yes | no |
| **Total** | 406 | 22196 |  |  |  |

**Table S4.** Estimated consensus genetic length by chromosomal recombination rate domains of Rockman and Kruglyak (2009). P-values are obtained by comparing the observed log change in genetic distance to a null distribution of log change obtained from 1,000 permuted datasets.

| Chr | Domain | Start (kb) | End (kb) | Size (kb) | Wild type size (cM) | rec-1 mutant size (cM) | P-value |
| --- | --- | --- | --- | --- | --- | --- | --- |
| I | Left arm | 855 | 2790 | 1935 | 20.6 | 21.9 | 0.383 |
|  | Center | 4636 | 8729 | 4093 | 4.8 | 32.7 | < 0.001 |
|  | Right arm | 11745 | 14467 | 2722 | 43.9 | 8.4 | < 0.001 |
| II | Left arm | 467 | 4392 | 3925 | 45.5 | 14.4 | < 0.001 |
|  | Center | 5815 | 11519 | 5704 | 7.5 | 38.9 | < 0.001 |
|  | Right arm | 12450 | 14911 | 2461 | 52.5 | 29.6 | 0.012 |
| III | Left arm | 670 | 3485 | 2815 | 44.9 | 13.9 | < 0.001 |
|  | Center | 4189 | 10009 | 5820 | 6.2 | 43.7 | < 0.001 |
|  | Right arm | 10745 | 13169 | 2424 | 41.6 | 27.8 | 0.043 |
| IV | Left arm | 1278 | 3355 | 2077 | 64.6 | 20.8 | < 0.001 |
|  | Center | 4815 | 12909 | 8094 | 25.5 | 50.0 | < 0.001 |
|  | Right arm | 13532 | 16875 | 3343 | 17.0 | 15.8 | 0.376 |
| V | Left arm | 1349 | 3272 | 1923 | 14.6 | 17.2 | 0.303 |
|  | Center | 5395 | 16471 | 11076 | 33.9 | 57.3 | 0.005 |
|  | Right arm | 16607 | 19939 | 3332 | 41.4 | 6.2 | < 0.001 |
| X | Left arm (+ 6.3% of the center) | 1503 | 5958 | 4455 | 16.3 | 20.6 | 0.297 |
|  | Center | 5958 | 13184 | 7226 | 33.6 | 36.5 | 0.368 |
|  | Right arm | 14139 | 16710 | 2571 | 17.1 | 5.1 | 0.003 |

**Table S5.** For each chromosome tests of heterogeneity of recombination rates. P-values are shown for a difference between the *rec-1* wild-type and mutant alleles, as assessed with null distributions of Gini differences from 1,000 permuted datasets. P-value for the difference from homogeneous recombination obtained from 1,000 simulated datasets with homogeneous recombination landscape (see Methods).

| Chrom | Wild-type Gini | Mutant Gini | Homogenous Gini (simulated) | Wild-type/mutant P-Value | Homogenous/mutant P-Value |
| --- | --- | --- | --- | --- | --- |
| I | 0.50 | 0.41 | 0.28 | 0.208 | 0.003 |
| II | 0.65 | 0.39 | 0.27 | < 0.001 | 0.018 |
| III | 0.61 | 0.35 | 0.35 | < 0.001 | 0.475 |
| IV | 0.60 | 0.24 | 0.29 | < 0.001 | 0.87 |
| V | 0.53 | 0.32 | 0.25 | 0.002 | 0.042 |
| X | 0.31 | 0.34 | 0.19 | 0.84 | < 0.001 |

**Table S6.** Comparison of X chromosome non-disjunction rates between studies and strains.

| Reference | Strain | rec-1 allele | Males in selfed progeny % (n) | Temperature °C |
| --- | --- | --- | --- | --- |
| Rattray and Rose (1988) | N2 | wt | 0.18 (n = 1660) | 20 |
|  | BC313 | s180 | 0.39 (n = 4915) | 20 |
|  | N2 | wt | 0.88 (n = 1950) | 26 |
|  | BC313 | s180 | 1.7 (n = 1857) | 26 |
| Chung *et al.* (2015) | N2 | wt | 0.05 (n = 2574) | 20 |
|  | Unnamed | h2875 | 0.35 (n = 3118) | 20 |
| This study | EEV1401 | wt | 0.15 (n = 5308) | 20 |
|  | EEV1403 | eeg1403 | 0.48 (n = 4832) | 20 |
|  | EEV1402 | wt | 0.07 (n = 6082) | 20 |
|  | EEV1404 | eeg1404 | 0.28 (n = 5974) | 20 |

## Supplementary Figures

**Figure S1.**
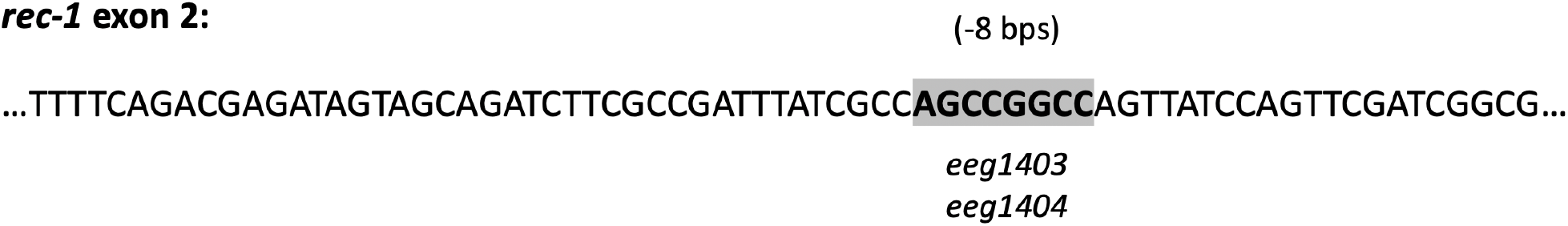
*rec-1* loss of function mutant obtained CRISPR-Cas9 directed mutagenesis of an 8 base pairs deletion in exon 2, leading to a truncated protein (Chung *et al*. 2015).

**Figure S2.**
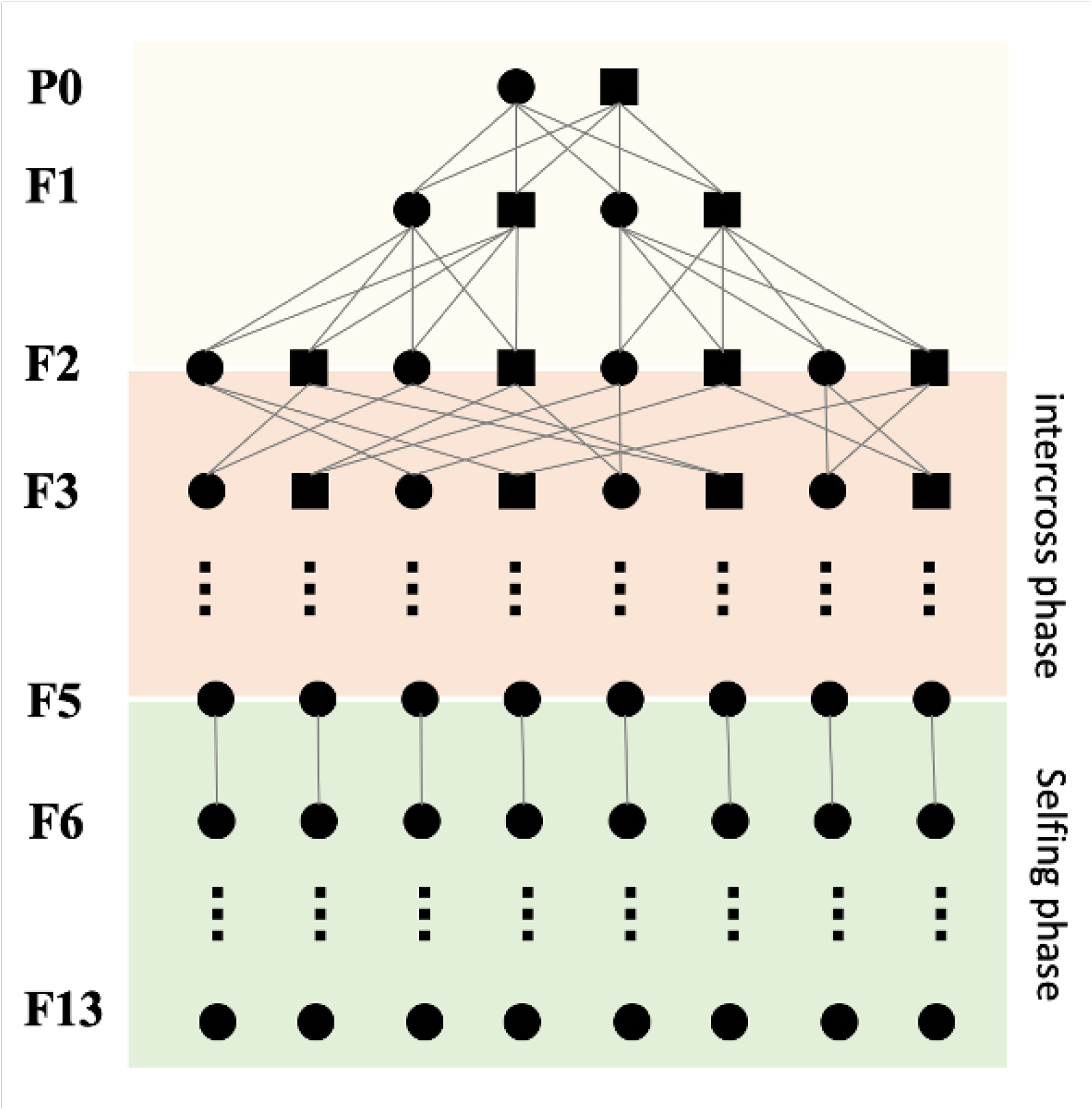
RPMEC-RIAIL construction. Illustration of the design employed for subpanel RIAIL construction. A given subpanel is initiated by mating one hermaphrodite (circle) and males (circle) from two different strains (*P*_0_) (see Table S2). Only one male is represented for convenience. *F*_1_ males and hermaphrodite are sib-mated and then 55 *F*_2_ hermaphrodite with two *F*_2_ males each. After that, until *F*_5_, a cross is set up between one hermaphrodite progeny of each cross and mated with two males from another cross during the intercross phase (orange). Each cross progeny contributes equally to the next generation with one hermaphrodite and two males. Following the intercross phase, one inbred line from each cross is derived for 8 generations by single hermaphrodite selfing (green).

**Figure S3.**
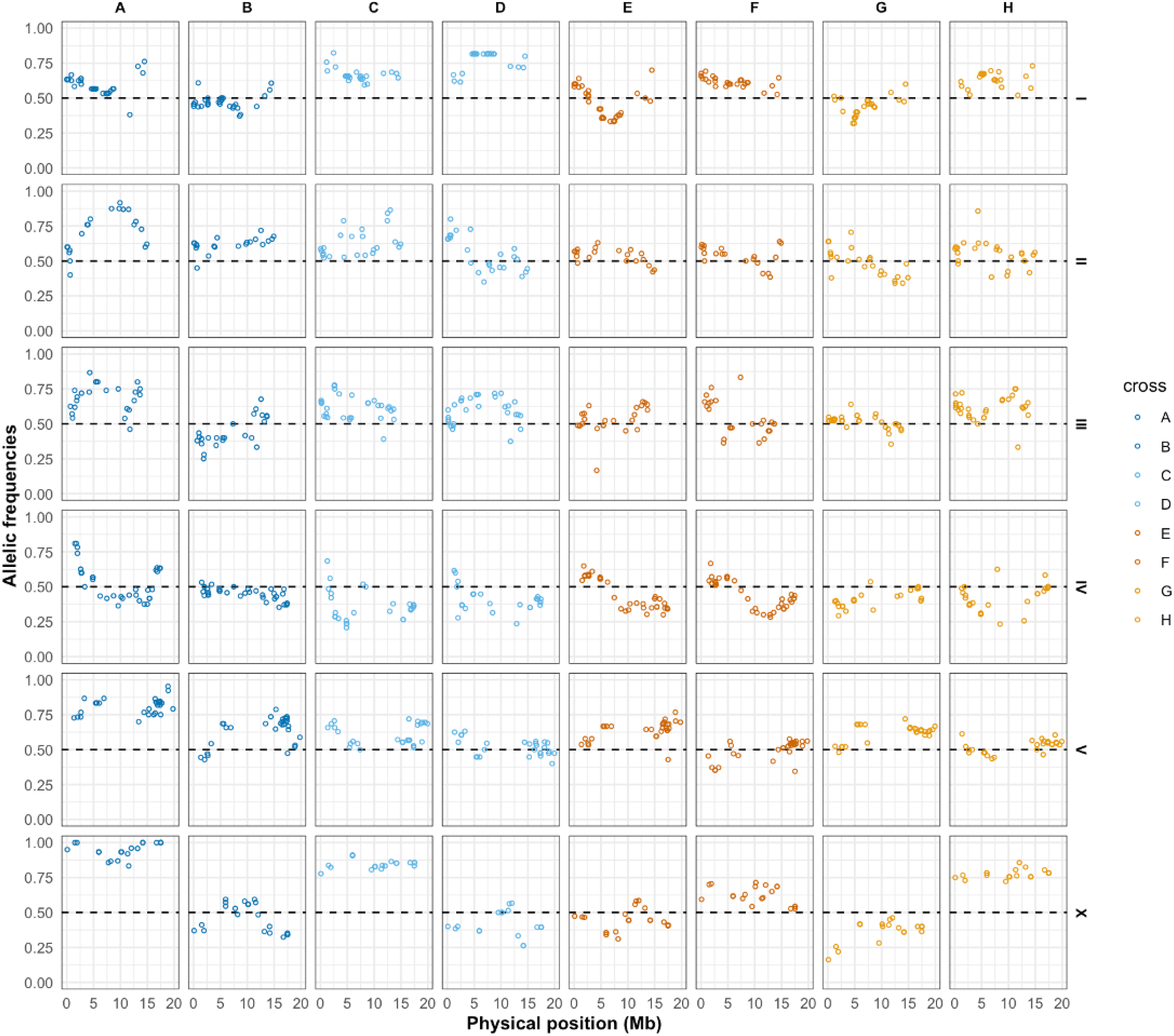
Allelic frequency deviation among intercross subpanels. Allele frequencies of EEV1401-polarized in *rec-1* wild-type (blue) and mutant (orange). Dark colors: intercross type 1; light colors: intercross type 2, including reciprocal crosses. See Table S2 for details about the A-H intercross subpanel types. The three highest mean allele frequency deviations from 50% are found on chromosome V subpanel A, and chromosome X subpanels A and B. The exclusion of these subpanels from the analysis has little effect on chromosome V estimates but may impact X chromosome estimates (see Figure S5).

**Figure S4.**
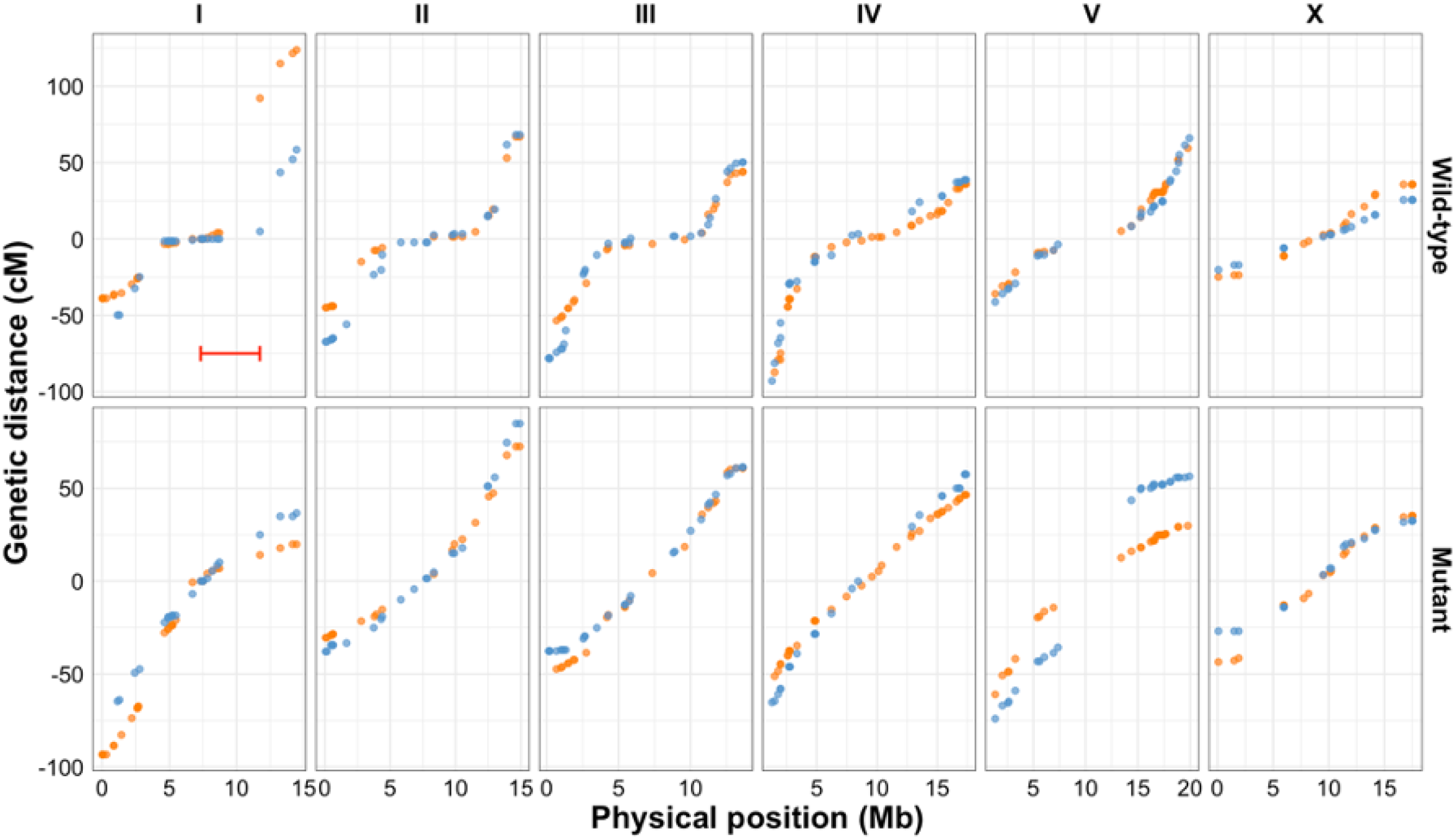
Intercross-specific Marey plots. Genetic linkage maps were estimated for the two intercross type panels differing in P0 parents’ genotype (orange: intercross type 1, blue: intercross type 2; see Table S2 for details). The estimated genetic distances of SNV markers are shown against their physical position. For each *rec-1* allele, we tested for significantly different genetic intervals (see main text for details). Only one interval in chromosome I is different between wild-type intercrosses (red bar).

**Figure S5.**
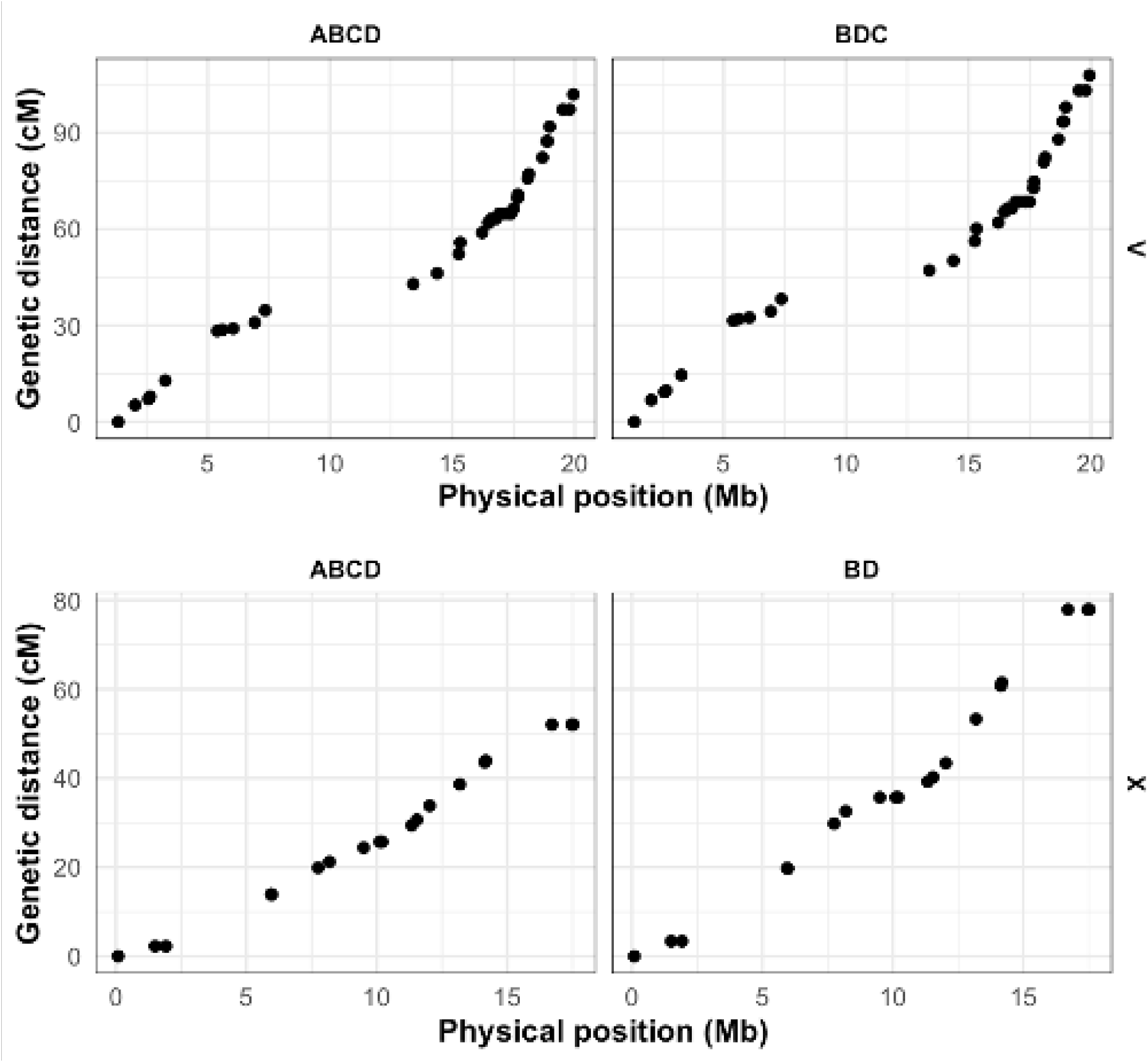
Withdrawal of subpanels with high SNV allelic deviations after RIAIL construction. The three highest mean allele frequency deviations from 50% are found on chromosome V in subpanel A (top), and chromosome X in subpanel A and B (bottom; see also Figure S3). Genetic linkage maps including (left) or excluding (right) the relevant subpanels.

**Figure S6.**
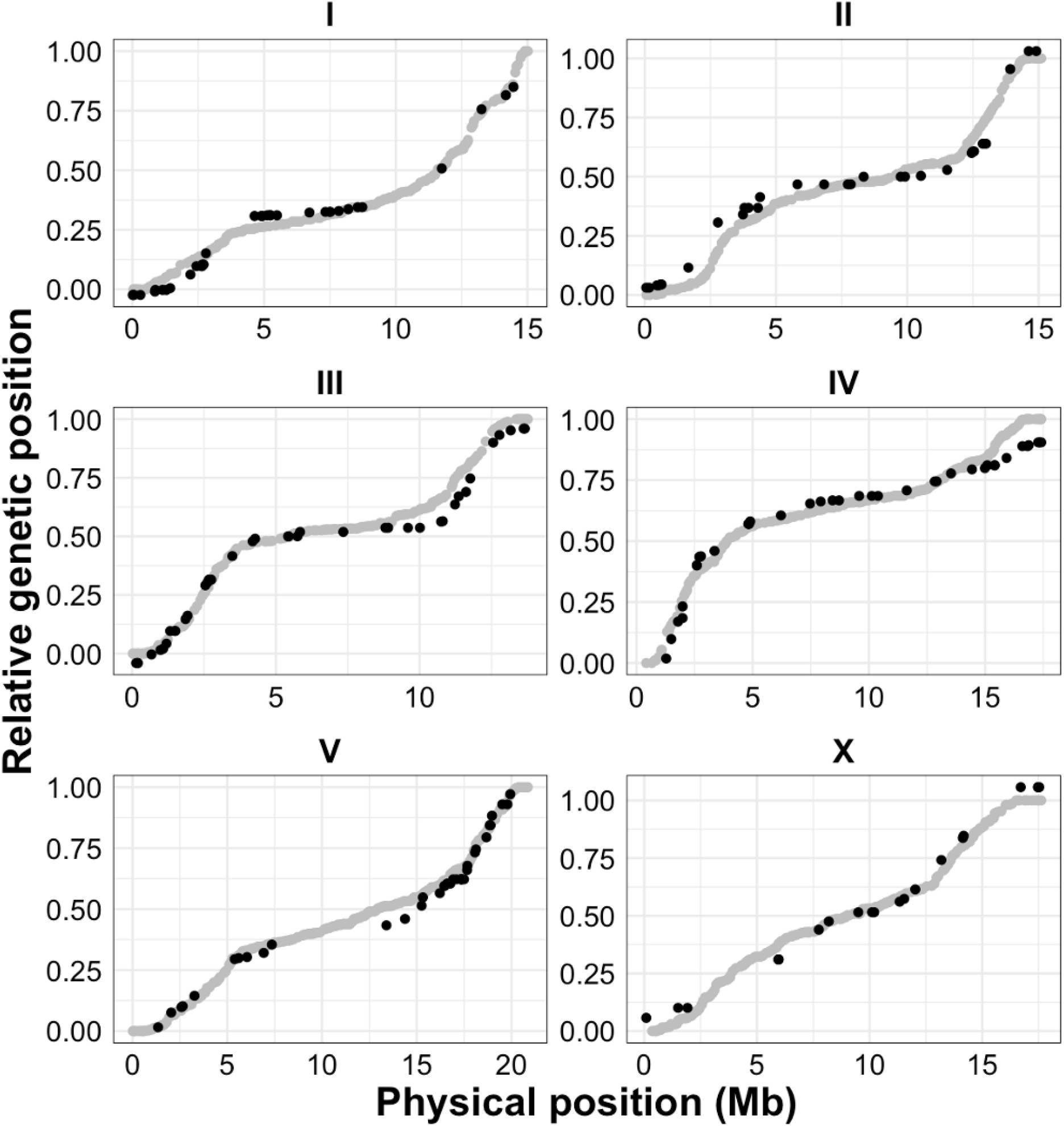
Comparison with the reference genetic linkage map. Marey plots showing relative genetic position as a function of the relative physical position for SNV markers (dots) measured in this study and for SNV markers reported by Rockman and Kruglyak (2009). Because the markers reported in this study did not span the entirety of all chromosomes, the relative genetic distances were corrected by multiplying it by the relative genetic distance between the extreme markers of Rockman and Kruglyak (2009). The genetic positions were then centered on a relative physical position of 0.5. Overall, there is a high congruence between the two maps. Interestingly, chromosome I show depressed recombination rates near the left subtelomeric tip, which was not observed by Rockman and Kruglyak (2009) but is consistent with the increased recombination rate in the *rec-1* mutant observed by Rattray and Rose (1988), and also observed here. Genomic incompatibility resulting from the *zeel1-peel1* system (Seidel *et al*. 2008) may explain map distortion in this region of Rockman and Kruglyak (2009). The *R* code this figure can be found in our [GitHub repository](https://github.com/ExpEvolWormLab/Paree_map/tree/main/maps).

**Figure S7.**
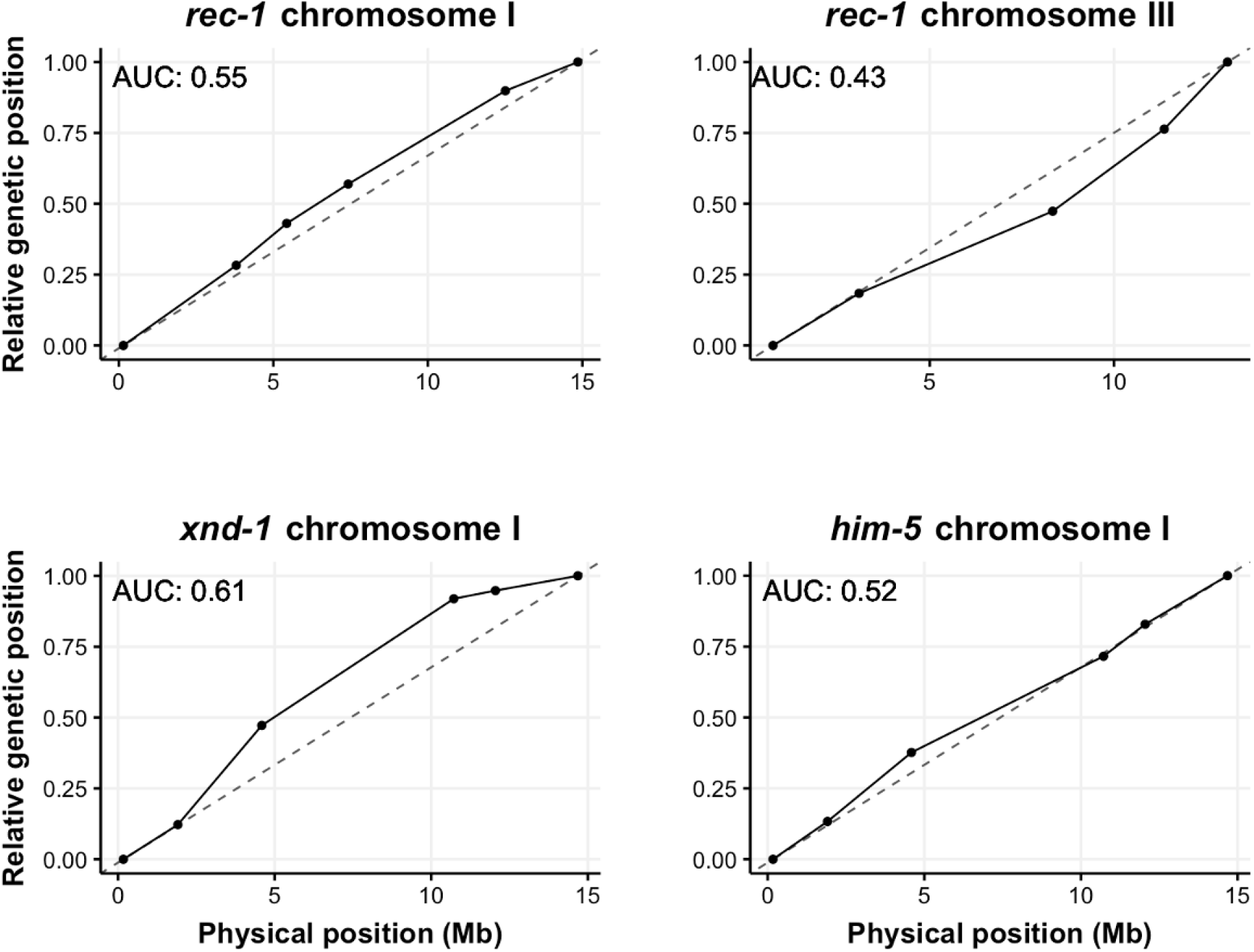
Curvature of published recombination landscape mutants. Top panels, recombination probability between a few morphological/molecular markers was estimated in a *rec-1* N2 strain mutant line for chromosome I by Zetka and Rose (1995) and for chromosome III by Chung *et al*. (2015). We show their relative genetic position against their physical position. Similarly, for two mutants in genes known to affect recombination rate landscapes: *xnd-1* from Wagner *et al*. (2010) and *him-5* from Meneely *et al*. (2012) for chromosome I. Area Under the Curve (AUC) is broadly similar to the estimates presented in Figure 4

**Figure S8.**
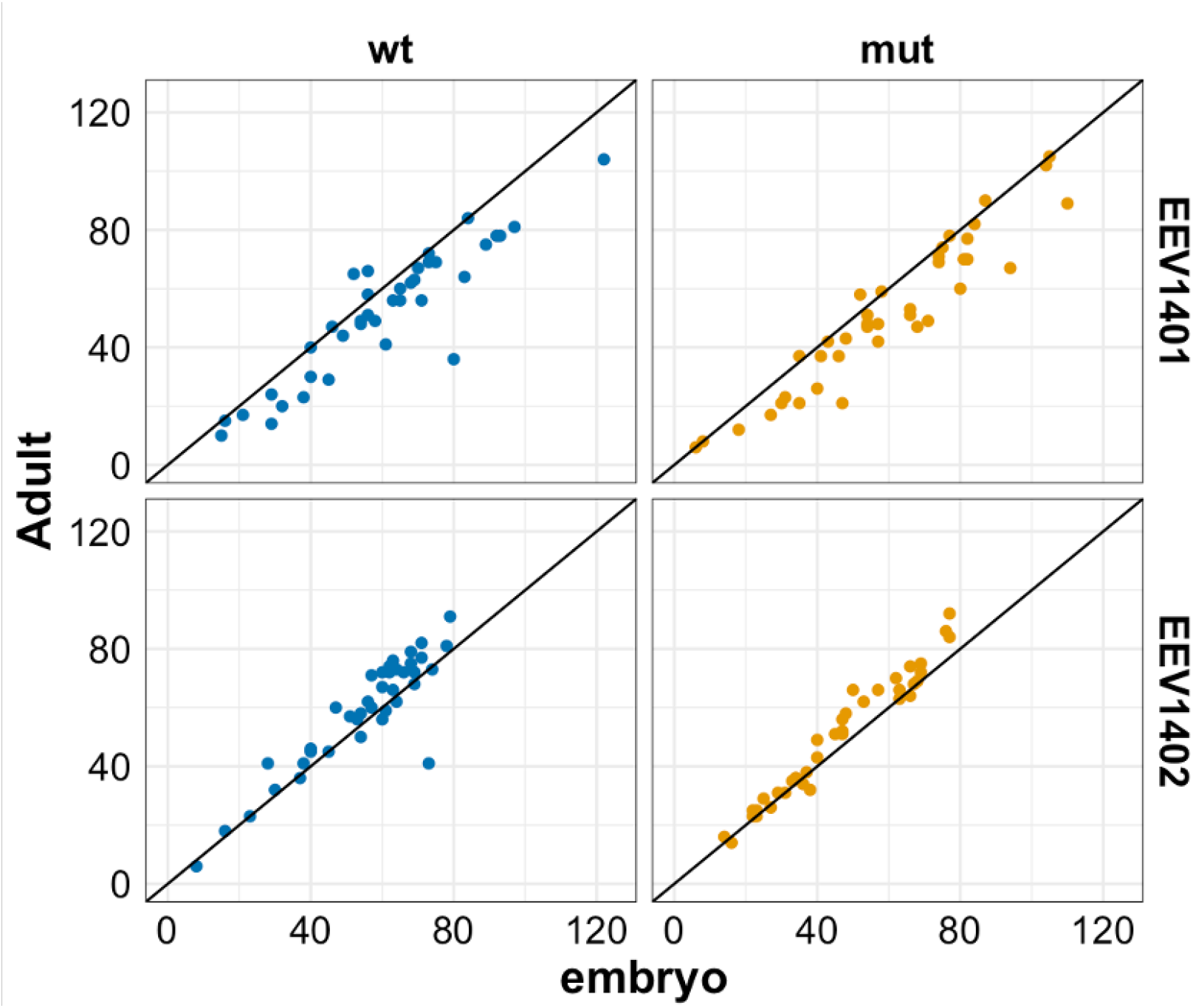
Hatchability differences between strains. Number of adults is shown over the number of embryos (dots) for two parental strains. Dots above the identity lines show that more adults than embryos were counted. There are no differences between the two rec-1 alleles (GLM; P-value EEV1401: 0.69; P-value EEV1402: 0.42). However, the EEV1401 genetic background has lower hatchability than the EEV1402 genetic background (GLM: p= 0.034)

